# Impacts of landscape-scale windthrow and subsequent, variable reforestation on bird communities in Central Europe

**DOI:** 10.1101/2020.06.18.159400

**Authors:** Johannes Kamp, Johanna Trappe, Luca Dübbers, Stephanie Funke

**Affiliations:** Institute of Landscape Ecology, University of Muenster, Heisenbergstr. 2, 48149 Muenster, Germany; Dachverband Deutscher Avifaunisten (DDA), An den Speichern 2, 48157 Muenster, Germany

**Keywords:** Extreme event, blowdown, vegetation succession, Norway spruce, farmland bird, long-distance migrant

## Abstract

With climate change, the area affected by and the intensity of forest disturbances such as windthrow, insect outbreaks and fire will be increasing. Post-disturbance forest management will be varied, and it is difficult to predict how much natural succession will be allowed in comparison to reforestation. Both, disturbance and reforestation will affect forest biodiversity globally, but potential shifts in species distribution, abundance and community composition are poorly understood.

We studied the response of breeding bird communities to windthrow and different reforestation strategies in one of Central Europe’s largest contiguous windthrow areas created by storm Kyrill in 2007. A decade after the disturbance, we compared bird species diversity, population densities and community composition on plots in replanted beech, replanted conifers and secondary succession (all salvage-logged after the storm), with undisturbed old Norway spruce *Picea abies* as a control, in the setting of a natural experiment.

Of the stands blown down, 95% were Norway Spruce. Reforestation strategies varied, with Spruce and non-native conifers planted on twice the area that was replanted with European Beech *Fagus sylvestris*. Large areas were still dominated by successional tree species a decade after the storm, especially birch, mirroring recommendations of sub-national forestry agencies to include secondary succession in future forest development. Birds responded strongly to windthrow, with a pronounced community turnover. Species associated with high conifer stands reached significantly lower densities on sample plots in disturbed areas. Replanted areas were characterized by mostly ubiquitous bird species. Areas dominated by secondary succession, especially birch *Betula spp*., were characterized by high densities of long-distance migrants (often species of conservation concern) and shrubland species, among them several indicator species.

Our results suggest that an increase of forest disturbance across Central Europe will lead to a pronounced reorganisation of biodiversity. Strategies that allow more secondary succession, and avoid replanting allochthonous tree species are likely to benefit populations of depleted bird species, even at salvage-logged and cleared disturbance sites.

## Introduction

Forest disturbance, i.e. large-scale tree loss due to mortality and the opening of the canopy, fundamentally affects ecosystem structure, properties and biodiversity in temperate and boreal forests (McDowell et al. 2020). Natural disturbance agents such as windthrow, insect calamities and fire (Schelhaas et al. 2003, Seidl et al. 2011) and disturbance by human activity affect 0.15% of the European forests annually, leading to billion m^3^ of wood loss (Schelhaas et al. 2003). Between 1986 and 2016, 17% of Europe’s forest area was disturbed, and disturbance frequency has been increasing consistently since 1986 (Senf & Seidl 2020).

With climate change, an increase in disturbed area and disturbance severity is expected globally, leading to younger forests with shorter trees (McDowell et al. 2020). Drought decreases the vitality of many tree species and can lead to die-backs (Steinkamp & Hickler 2015, Schuldt et al. 2020). In Central Europe, the area affected by forest canopy mortality has doubled over the past three decades (Senf et al. 2018, 2020). Apart from rising temperatures (Allen et al. 2010), extreme weather events such as severe storms are thought to increase in frequency globally as a result of climate change (Rahmstorf & Coumou 2011, Trenberth et al. 2015). This will affect European forests directly (Lindner et al. 2009, Seidl et al. 2017, Panferov et al. 2009), and will lead to more insect calamities that often radiate out from weakened stands after windthrow (Dobor et al. 2020).

A major disturbance agent in Central Europe are windstorms. Windstorm intensity and forest damage has increased between 1950 and 2010 across Western and Central Europe, with a further acceleration since 1990 (Gregow et al. 2017, Senf & Seidl 2020). The risk of windthrow is exacerbated by decreasing tree vitality due to climate-change induced stress, and insect calamities (Seidl et al. 2017). Canopy gaps resulting from windthrow allow new forest succession, thereby altering tree species composition and plant species richness patterns (Jonášová et al. 2010, Ulanova 2000). Windthrow leads to an increase in sunlight and windspeed at the soil surface, which can result in a decrease of relative humidity despite more available precipitation than in closed-canopy forest (Ishizuka et al. 2002). Disturbed sites are often characterized by increased soil nutrients contents (Ulanova 2000), and an increased habitat and micro-site diversity.

Across Central Europe, the resilience of forests to windthrow has decreased due to changing forest management over the past centuries. Native forests were dominated by deciduous tree species such as European Beech (*Fagus sylvatica*) and Oak (*Quercus spp*.). Since the late 18^th^ century, the area planted with coniferous trees like Norway Spruce *Picea abies* and Scots Pine *Pinus sylvestris* has strongly increased, and deciduous and mixed stands were transformed into uniform, dense conifer plantations across vast areas (Kirby & Watkins 2015). Less productive silvicultural systems such as wood pastures and coppice as well as degraded heathland were afforested with coniferous trees (McGrath et al. 2015, Plieninger et al. 2015). Spruce and pine were often preferred over deciduous tree species as they provide faster financial revenue. They also grow well on the degraded soils that resulted from intensive silvicultural use. Due to their root architecture and stability, coniferous trees are more susceptible to windthrow than closed stands of deciduous trees, and spruce is twice as susceptible as pine under comparable soil conditions (Ruel 1995). An increasing proportion of contiguous spruce stands in the landscape will therefore increase windthrow risk (Panferov et al. 2009).

The changed site conditions (microclimate, soil, vegetation, and structural heterogeneity) at windthrow sites have strong effects on biodiversity. Plant species richness is often higher at disturbed compared to undisturbed forests (von Oheimb et al. 2006). Windthrows in Europe act as regional hotspots for forest insect communities as they maintain open habitat continuity in a mosaic landscape and provide resources for saproxylic species, flower visiting insects and phytophages on saplings (Bouget & Duelli 2004). Other invertebrate communities can be affected negatively, e.g., earthworm species richness and biomass decreased with increasing disturbance intensity (Nachtergale et al. 2002). Bat species respond in various ways to forest gaps resulting from disturbance, depending on their wing morphology (Fukui et al. 2011, Mehr et al. 2012). Bird species richness can also increase after disturbance (Żmihorski 2010, Thorn et al. 2016), and abundance and community composition vary with species traits and across spatial scales (Murakami et al. 2008).

As a key management process after disturbance, the effects of salvage logging on biodiversity have received great interest over the past years (Thorn et al. 2018). However, most studies were conducted within 5 years after disturbance (Thorn et al. 2018), and few are from Europe (Georgiev et al. 2020). Furthermore, salvage logging will rather be the norm than the exception away from protected areas (Leverkus et al. 2018), as part of an intensive forestry management maximizing financial revenues. Studies comparing biodiversity responses to reforestation in commercially used areas are rare, especially those comparing different reforestation strategies (Żmihorski 2010). The lack of studies on biodiversity responses to alternative reforestation is unfortunate, as with accelerating climate change, reforestation strategies need to be developed that consider forest resilience, economic aspects, but also biodiversity conservation (Müller et al. 2019).

In Central Europe, a series of severe storms has created very large windthrow areas during the past 30 years (Senf & Seidl 2020). An exceptionally violent storm was ‘Kyrill’ on 18 and 19 January 2007. Kyrill caused damage estimated at 10 billion US$ across Western Europe, with 47 fatalities, power cuts and severe damage to transport infrastructure and buildings (Burghoff et al. 2010). In its wake, windthrow occurred over vast areas, with ca. 62 million trees uprooted or windsnapped, equal to 59 million m^3^ of standing wood damage (Fink et al. 2007). Exceptionally large windthrow areas are found in Germany (Fink et al. 2007, Wald & Holz NRW 2019). These were salvage-logged within one or a few years after the storm, with a few exceptions such as in protected areas (Thorn et al. 2016). Despite salvage logging, due to the sheer number and size of windthrows, not all have been reforested a decade after the storm, and areas of natural succession exist. Reforestation strategies range from “traditional”, dense spruce plantations to mixed stands and reforestation with beech and other deciduous species. In Germany, most of the Kyrill windthrow affected private forests that are often small in size and fragmented. The variation in windthrow size and reforestation strategy provides an ideal “natural experiment” for exploring biodiversity development after windthrow in a real-world, landscape-level context.

We studied the response of biodiversity to different reforestation strategies a decade after windthrow in a space-for-time design. We used birds as they are good indicators of overall biodiversity, and ecological information e.g. on functional traits is readily available. We aimed to provide baseline knowledge to inform future reforestation strategies, considering that forest disturbance events are likely to become more frequent in the near future due to climate change.

Our objectives were:

1. To map and quantify the area of different reforestation strategies in a landscape characterized by very large windthrow areas.
2. To explore shifts in bird abundance and community composition after windthrow by comparing blowdowns to remaining, old spruce stands a decade after disturbance.
3. To predict differences in bird abundance and community composition as a function of different reforestation strategies.

## Methods

### Study area

We conducted a field study in the “epicentre” of Kyrill windthrow (Fink et al. 2007) in Germany, in the district Märkischer Kreis of Northrhine-Westfalia. Here, ca. 50,000 ha Norway Spruce were blown down by Kyrill in a single night, with very large contiguous patches of up to 211 ha (Wald & Holz NRW 2019) that were all salvage-logged after the storm. The study area of 1062 ha (ca. 902 ha forested before Kyrill, Fig. 1, Fig. 2) is located in the Balve forestry department east of the village of Garbeck (51.3175°N, 7.829167°E), a city district of Balve. The area is part of the geographic region “Sauerland” and has a humid climate (annual precipitation sum 923 mm, annual mean temperature 8.2°C). Altitude in the surveyed areas ranges between 279 m and 441 m a.s.l. The area is characterized by comparatively steep valleys and exposed hills, and vulnerable to windthrow due to this topography.

**Figure 1:**
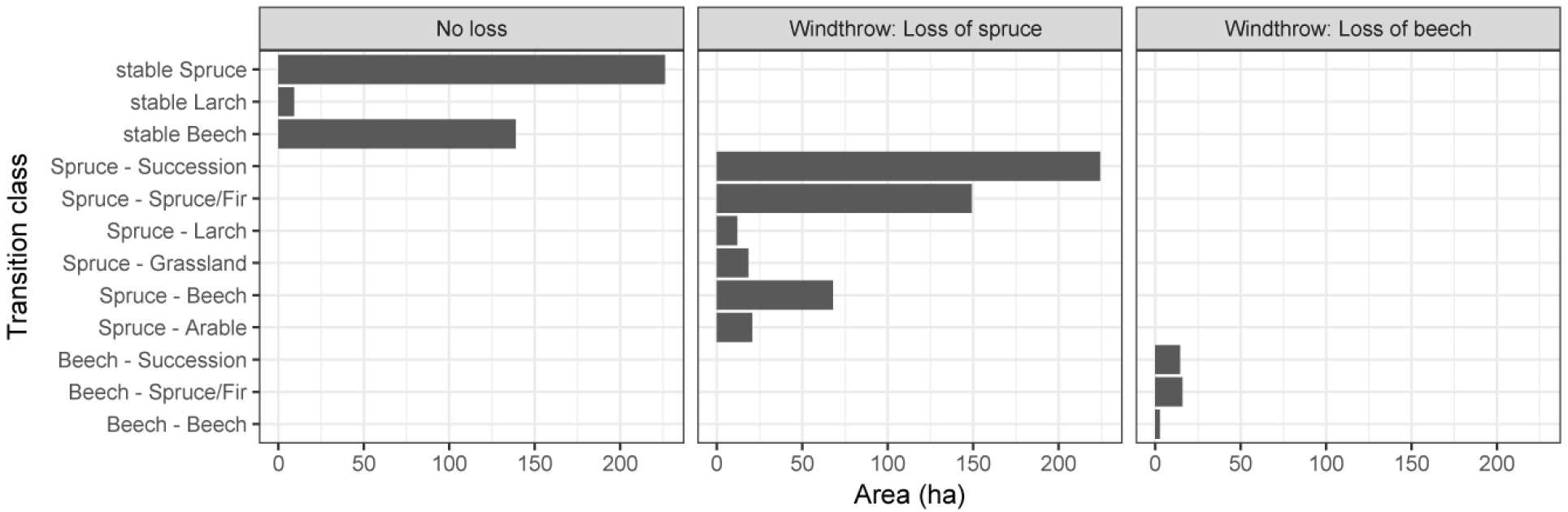
Management transitions between habitat types in the study areas comparing the period before windthrow (2007) with the situation in the year of the bird surveys (2018). “Succession” comprises areas still dominated by secondary succession (mostly Birch). Some of these have not seen interventions by forestry, but in some, successional trees have already been girdled and beech has been planted under the canopy.

**Figure 2:**
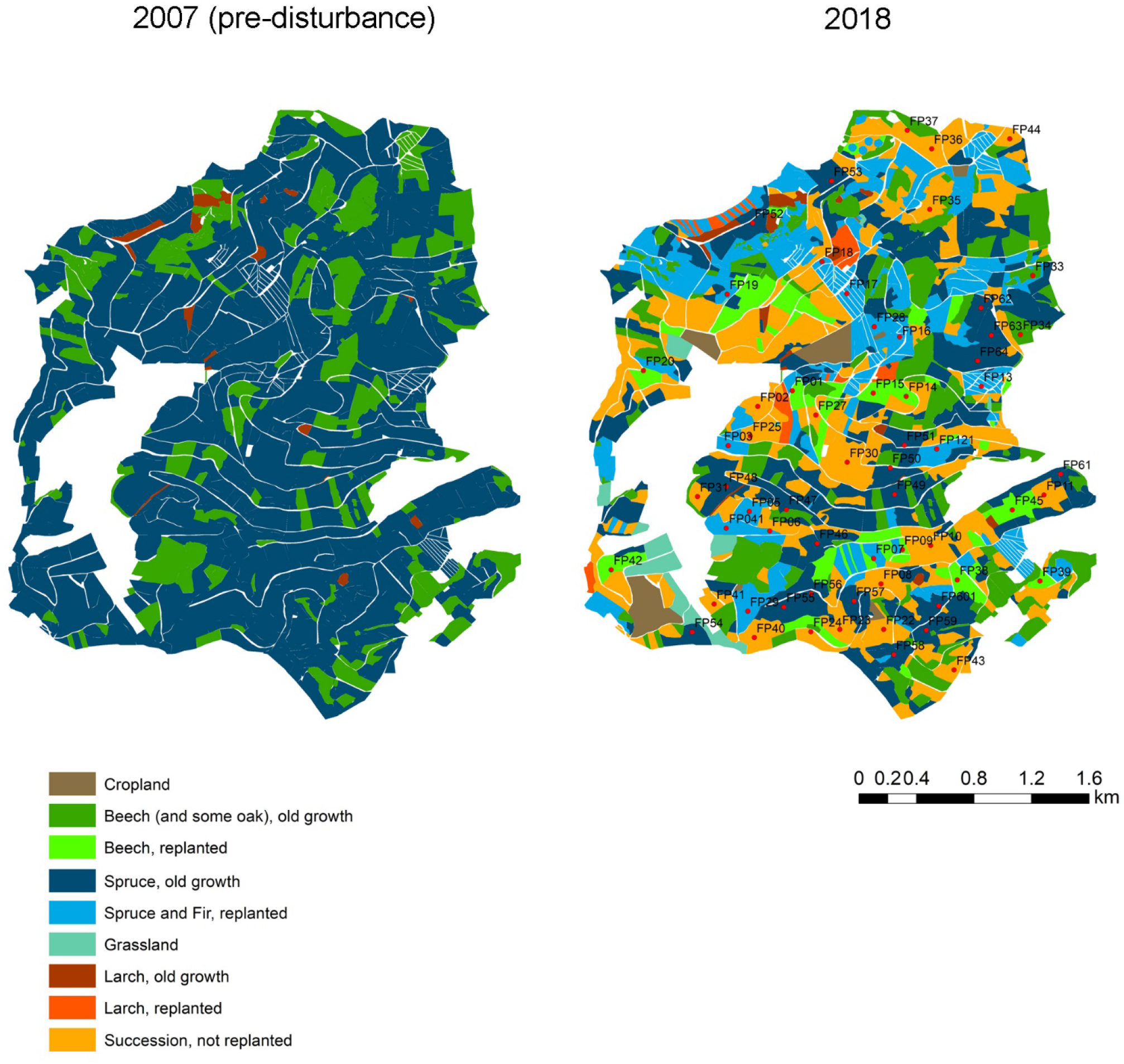
Pre-disturbance forest habitats in 2007, before the windthrow event, and in 2018 after reforestation. Categories contain the trees that dominated the respective area. Bird sampling plots are given as red dots on the 2018 map, plot names correspond to those in the raw data file.

### Mapping windthrow and reforestation

We were interested to quantify the spatial distribution of windthrow and areas of different subsequent reforestation strategies to develop a spatial basis for the bird sampling design. Official shapefiles of Kyrill windthrow areas from Opengeodata NRW (2020) and forest inventory maps were used to pre-select areas disturbed in 2007 to obtain a map of available windthrow areas, but both sources were rather coarse and did not allow to unambiguously map reforestation strategies and changes in land use following the storm event. We therefore manually digitized current and pre-disturbance forest types with more detail from a time series of cloud-free, ortho-rectified aerial images with 0.4 m resolution (examples in supplementary online material, Text S1). These were provided by the German Federal Agency for Cartography and Geodesy and are publicly accessible via Google Earth (copyright label: GeoBasis-DE/BKG). Windthrow areas were mapped from an image taken on 31 March 2009 (Text S1, Fig. S1). At this time, all windthrow areas had been cleared of trunks and salvage logging was completed, but the areas had not re-vegetated yet. (We did not detect any areas that were not salvage-logged and cleared from tress.) The root plates of uprooted trees were still visible, as were harvester tracks and piles of wood near forest tracks. The imagery allowed distinguishing between windthrow of Norway Spruce (*Picea abies*) stands (smaller, regularly spaced root plates and grey soil surface coloration) vs. European Beech (*Fagus sylvatica*) stands (larger, irregularly spaced root plates and pale brown soil surface). The classification of spruce vs. beech blowdowns was double-checked with a Landsat 7 ETM image from summer 2006, where spruce stands appeared dark green, whereas deciduous stands were pale green. We could not distinguish between pre-disturbance Norway Spruce and Douglas Fir (*Pseudotsuga menziesii*) stands. Douglas Fir is occasionally cultivated in the area, but comprises only 2% of all trees in German forests (Federal Ministry of Food and Agriculture 2015). Forestry maps suggested that some of the stands perceived as European Beech also contained old oaks (*Quercus* spp.), but field visits revealed that beech dominated by far.

The extent and distribution of reforestation at the time of our study were mapped on two further aerial images dated 27^th^ March 2017 (before leaf emergence on deciduous trees, Text S1, Fig. S3) and 4^th^ May 2018 (after emergence, Text S1, Fig. S4), corresponding with the start of our field survey. We distinguished between secondary succession (areas with natural regrowth of trees of early successional stages, mostly European White Birch *Betula pendula*), and tree plantations (henceforth reforestations). The latter consisted of conifers or European beech (see below, ground truthing). On the March image, areas of secondary succession appeared as a mix of irregularly spaced deciduous trees and sparse conifers, whereas both conifer (spruce and fir, dark green) and beech plantations (brownish, since leaf-less) were easy to detect by their regular spacing in planting rows and the small size of trees. Remaining, not thrown spruce stands were easy to map as single trees were distinguishable by shape, and irregular stand borders indicated windthrow. European Larch (*Larix decidua*, both older trees and reforestations) were mapped on the March image as they had a distinct conifer appearance, but were greyish compared to the dark green spruce and fir. The May image allowed to confirm the mapped areas of beech reforestation that appeared pale green, and areas of secondary succession that were dominated by the pioneer tree European White Birch (*Betula pendula*) and appeared darker green. Mature European Beech stands could easily be mapped from an image taken on 11^th^ October 2015 (Text S1, Fig. S2) due to the characteristic orange autumn leaf coloration. All polygons were delineated in google earth, saved as kml-files, imported into and further processed in Arc GIS 10.6.

To ground-truth the mapping of the reforestation areas, tree cover and height were measured at all 60 randomly selected bird plots (see below). During a further ground-truth survey in the north of the study area (that had less bird sample points than the southern part), we opportunistically visited 204 polygons and took notes on tree composition and cover. Therefore, a total of 244 (25.6%) of the 953 mapped polygons were ground-truthed in the field. Field surveys revealed that all areas mapped as “spontaneous regrowth” in our remote sensing approach were indeed still dominated by pioneer trees (especially birch), but that on a part of them forestry management had taken place that aimed at a gradual transition of the areas into commercially marketable wood resources. This included i) planting of spruce, fir and beech under the canopy of the pioneer trees, occasionally occupied by girdling the dominating birch trees, ii) thinning of the pioneer trees to make space for the planted trees. Field surveys also revealed considerable heterogeneity in conifer reforestations, which were partly also used for Christmas trees and the harvest of ornamental twigs (the latter leaving behind some areas of the reforestation without vertical branches).

### Bird surveys

We surveyed breeding birds in the field using point counts, combined with distance sampling (Buckland et al. 2015) as we wished to estimate bird densities. We compared bird densities across the three “treatments” of the natural experiment (beech reforestation, conifer reforestation, secondary succession) and the control (old spruce forest). We selected 80 points in a randomly stratified approach based on the 2018 habitat map (see above), with an initial 20 points in each habitat category. Twenty points had to be discarded, as they were not accessible due to property rights, or were in areas currently worked by forestry. This resulted in a final sample of 60 points available, 18 in old spruce forest, 12 in beech reforestation, 13 in conifer reforestation, and 17 in secondary succession (Fig. 1). During selection, points were constrained to be at least 200 m apart to minimize the risk of double counts, and to be 50 m away from the border of a habitat class of to avoid strong edge effects. Three repeat counts were conducted across the points between 03 April and 18 May (which is the peak activity period of breeding birds in Central Europa), leading to a total of 180 point counts available for analysis. Surveys were conducted on days with suitable weather conditions (no strong wind, rain or fog, no frost) from dawn until 11 a.m. by authors LD, SF and JK who are all experienced observers. Observer bias was limited by conducting joint trial counts and distance measurements in the beginning of the season. Birds were recorded in five distance bands: 0–10 m, 10–25 m, 25–50 m, 50–100 m, 100–200 m. Range finders were employed throughout to measure distances to birds or the presumed location of individuals detected through aural cues. Territorial and non-territorial birds were recorded separately. All bird counts lasted 5 minutes, with a 5-minute settling period. Flyovers were excluded, but not birds hunting in flight at the points. For analysis, only records of territorial birds were used, as 95.5% of the detections referred to territorial (mostly singing) birds. Taxonomy and nomenclature follow Handbook of the Birds of the World and BirdLife International (2019).

### Habitat data

To allow an identification of potential structural drivers of forest bird “successions” we recorded a number of vegetation parameters. We delineated a recording quadrat of 10 × 10 m centred on each survey point and measured vegetation height (with a folding meter, to the nearest cm) at the four corners. These four values were then averaged for analysis. We also visually estimated the cover of the following parameters: tree (>1.5 m height), shrub (0.5-1.5 m) and herb layer (<0.5 m height), beech, birch, spruce, Douglas fir, other successional tree and shrub species (mostly *Sorbus aucuparia*, *Sambucus nigra*, *Salix caprea* and *Alnus glutinosa*), *Rubus fruticosus agg*. shrub, dry plant litter, bare ground and woody debris. The vegetation structure surveys were conducted once, around the second survey round in early May when leaves had already unfolded, during the annual activity peak in most of the surveyed bird species.

### Data analysis

All statistical analyses were carried out in R 3.5.1 (R Core Team 2020). All data collected and analysed in this study are available as supplementary online material, Table S1.We were first interested to describe how bird abundance varied across the four habitat categories. We used a hierarchical distance sampling modelling framework for closed populations to estimate population densities, while accounting for varying detection probability and availability for detection of the study species (Kéry & Royle 2016). We modelled the density of territorial individuals, i.e. considered only singing males. From earlier explanatory analyses, we inferred that model fit was generally poor when the incidence of a species fell to <10% occupied sampling units. We therefore considered only species for distance sampling analysis that occurred at more than 10 out of the 60 surveyed points.

We fitted a two-part hierarchical distance sampling model. We considered the variables observer and vegetation height in the detection probability part of the models, as we assumed that detection probability might decay faster with distance when observers were less experienced, or vegetation taller and denser. However, there was little support for an effect of either variable (with significance of both variables consistently being p >0.05) so we included only the intercept in the detection part of the final models. A single categorical variable was included in the abundance part of the model, the habitat type (old spruce vs. different reforestation categories). The models therefore yielded a density estimate across all sample points per habitat type. All models were fitted using the ‘distsamp’ function in package unmarked (Fiske et al. 2011), using a Poisson model with a half-normal detection function. Estimates densities were considered significantly different where 95% parametric confidence intervals did not overlap.

For further analyses, we pooled the three visits by selecting the visit with the highest count for each plot-species-combination. These maximum values were then used in further analyses, and to calculate species richness and Shannon diversity for each of the 60 plots. To identify species associated with old spruce forest compared to those of the windthrow areas (reforested with beech or conifers, and secondary succession), we performed an Indicator Species Analysis in the R package indicspecies, De Caceres & Legendre 2009) permitting only single group affiliations. Indicator values are scaled between 0 and 1, and the index is maximum when all individuals of a species are found at a single group of plots and when the species occurs in all plots of that group. For each species, the best group association was tested for statistical significance. We were also interested to assess how bird community composition was affected by reforestation strategy, and the resulting vegetation structure. We explored community composition in 2018 using non-metric multidimensional scaling (NMDS), a common ordination technique. Through ordination, sites can be visualized in multidimensional space according to their similarity in species composition. Likewise, species can be displayed according to their similarity in occurrence at the sample sites. The NMDS was based on the Bray-Curtis dissimilarity index and performed with the function ‘metaMDS’ from the R package vegan (Oksanen et al. 2018). The ‘envfit’ function allowed us to then fit centered and scaled (z-transformed) environmental factors onto the ordination.

Finally, we sought to gain insight in species community change after windthrow by examining groupwise responses of functional “guilds”, that is species groups sharing similar ecological and functional traits (Thorn et al. 2016). Species were assigned to guilds according to their migratory behaviour (long-distance, short-distance, resident), nest position (closed, hollow, niche, open), diet (omnivore, invertebrates, vertebrates, fruits/seeds) and feeding site (ground, vegetation, trunk, indifferent). Trait information was obtained from a Germany-wide database (Wahl et al. 2015, unpublished) and Thorn et al. (2016) (supplementary online material, Table S2). For each guild, the abundance of individuals per plot was calculated from the raw data (no distance sampling densities were used as these could not be derived for all species, see above). Generalized linear models (GLM) were then fitted with guild abundance as response variable and habitat type as the explanatory variable. For all models but open nest breeders (normal distribution) we used Poisson distributions with a log link. To disentangle trends resulting from overall abundance, the total number of individuals per plot was set as an offset term. All GLMs were fitted using the ‘glm’ function from the package stats (R Core Team 2020) and model fit was assessed with functions implemented in the DHARMa package (Hartig 2019). Based on the models, estimated marginal means (EMMs) for each habitat type as well as contrasts between habitat types were calculated using ‘emmeans’ from the emmeans package (Lenth 2019).

## Results

### Forest change

In total, 58.8% of the forests in the study area were lost through windthrow. Windthrow affected mostly spruce plantations (68% of the 2006 area thrown, Fig. 1, Fig. 2), deciduous trees were affected less (20% of the 2006 area thrown, mostly beech, Fig. 1). Spruce represented 94.4 % of the thrown trees. To a small extent, windthrow areas were converted to cropfields and pasture (7.4%). Reforestation with significant intervention between 2007 and 2018 (clearing the area, soil treatment, row planting of trees) was detected on 13.5% of the windthrow area (beech) and 33.4% of the area (conifers, mostly spruce and Douglas fir, Fig. 1). Areas dominated by successional trees (but partly with some forestry intervention and replanting under canopy, see above) still covered 45.4% of the area in 2018.

### Species richness, diversity and abundance

Forty-nine species were recorded across all sample points, ranging from 7 to 20 (mean 12.4) species per plot. There was no difference in overall species richness (Kruskal-Wallis-test, χ^2^_46_ = 1.630, *P* = 0.653), Shannon diversity (Kruskal-Wallis-test, χ^2^_46_ = 1.007, *P* = 0.800) or total abundance (Kruskal-Wallis-test, χ^2^_46_ = 0.533, *P* = 0.912) between the four habitat types.

Twenty-one species occurred at more than 10 out of the 60 surveyed points and were thus available for abundance estimation with distance sampling methods (supporting online material, Table S3). Of these species, eight were “disturbance winners”, i.e. showed significantly higher densities of territorial males in at least one disturbed habitat category compared to the baseline (old spruce; Fig. 3). Willow Warbler, Garden Warbler, Tree Pipit and Yellowhammer appeared only in disturbed habitat (with the exception of a single Willow Warbler), whereas Dunnock had significantly higher densities in conifer reforestation, and Common Chiffchaff, Eurasian Blackcap and European Robin in beech reforestation suggesting preferences for these reforestation habitats.

**Figure 3:**
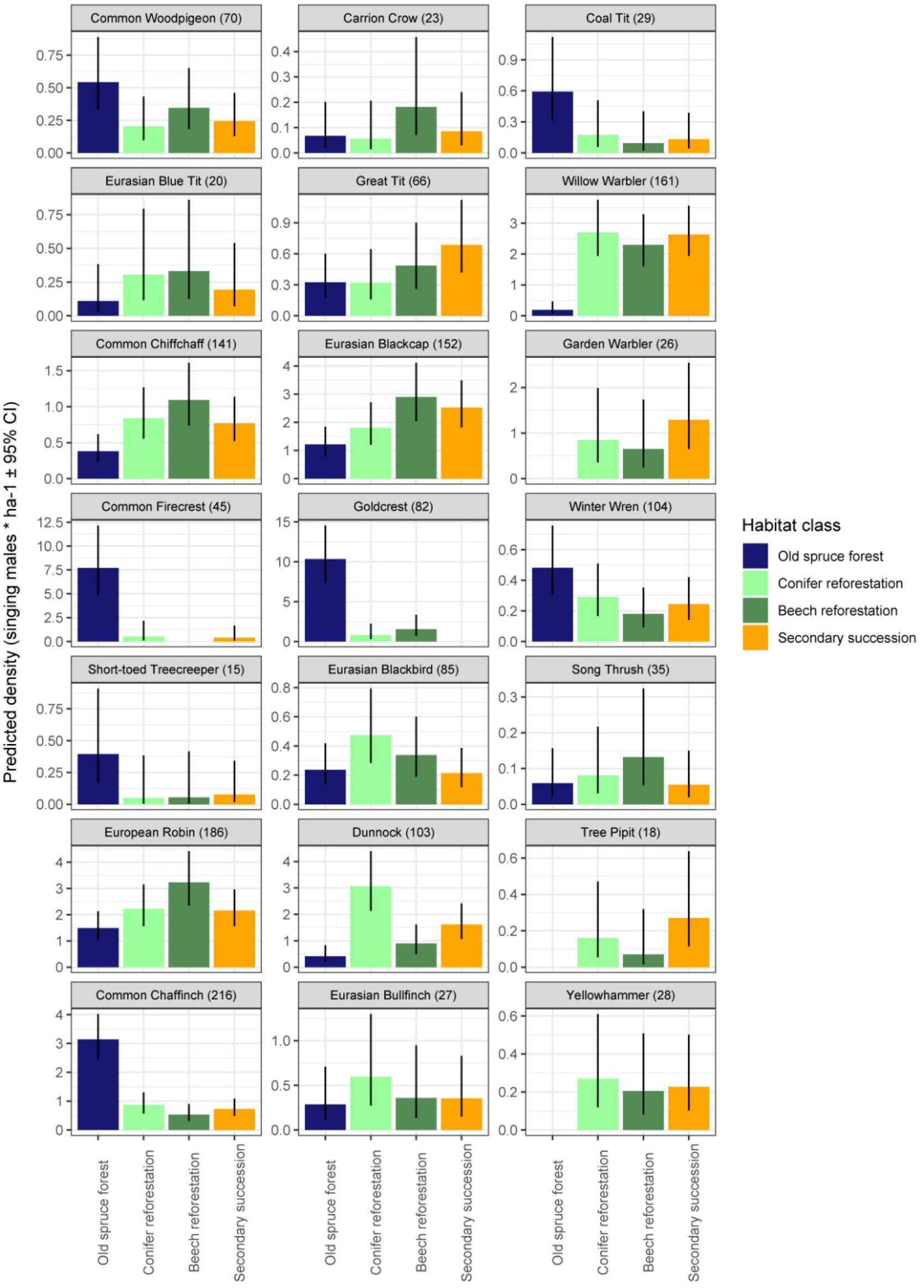
Predicted density of singing (=territorial) males as a function of habitat and reforestation strategy from hierarchical distance sampling models, for the 21 most common species in 2018. Numbers in brackets give total sample size (territorial individuals summed over all three counts).

Six species were “disturbance losers”, i.e. showed lower densities in all disturbed habitat category compared to the old spruce (significantly lower in Common Firecrest, Goldcrest and Common Chaffinch; Fig. 3). The remaining seven species showed not significant pattern in abundance differences across habitat categories (Fig. 3). Estimated densities were in the expected order of magnitude suggesting that distance sampling assumptions were met. Exceptions were Common Firecrest and Goldcrest. Densities for these were at least threefold higher than reported in the literature suggesting that a consistent underestimation of distance to observer led to a steep decline of the density function after the first distance category of 10 meters. This is unsurprising as the low-volume, but high-pitched voices of both species reach further than observers commonly assume.

### Indicator species

Indicator species of remaining old spruce stands comprised a typical set of species that colonized large parts of Central Europe in the wake of the extensive conifer plantations of the 19^th^ century, such as Common Firecrest and Goldcrest, European Crested Tit and Coal Tit (Table 1). Indicator species of reforested areas comprised few high-forest species. Reforestations were mainly characterized by ubiquitous, common species. Indicator species of secondary succession included a number of declining long-distance migrants, such as Willow Warbler, Garden Warbler, Tree Pipit and Common Whitethroat (Table 1).

**Table 1:**
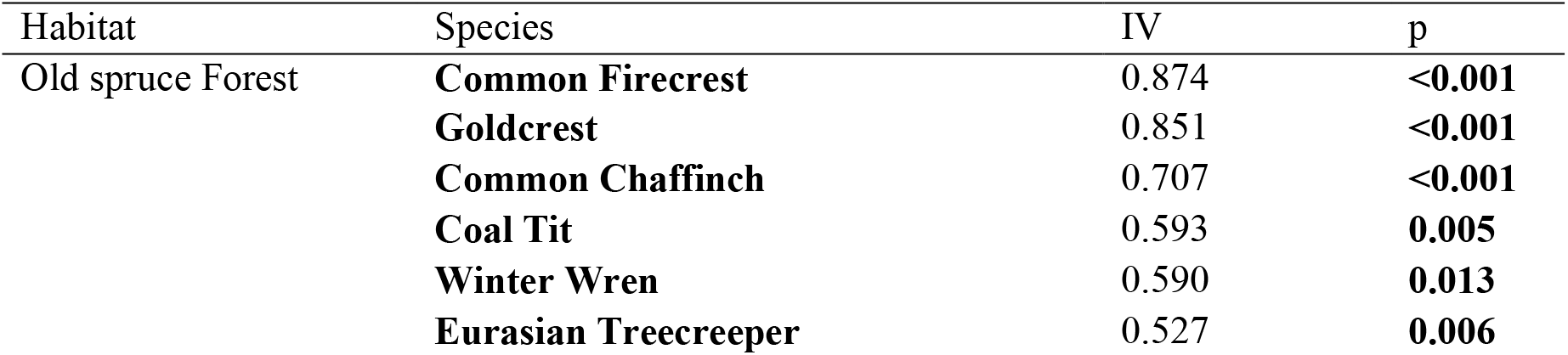
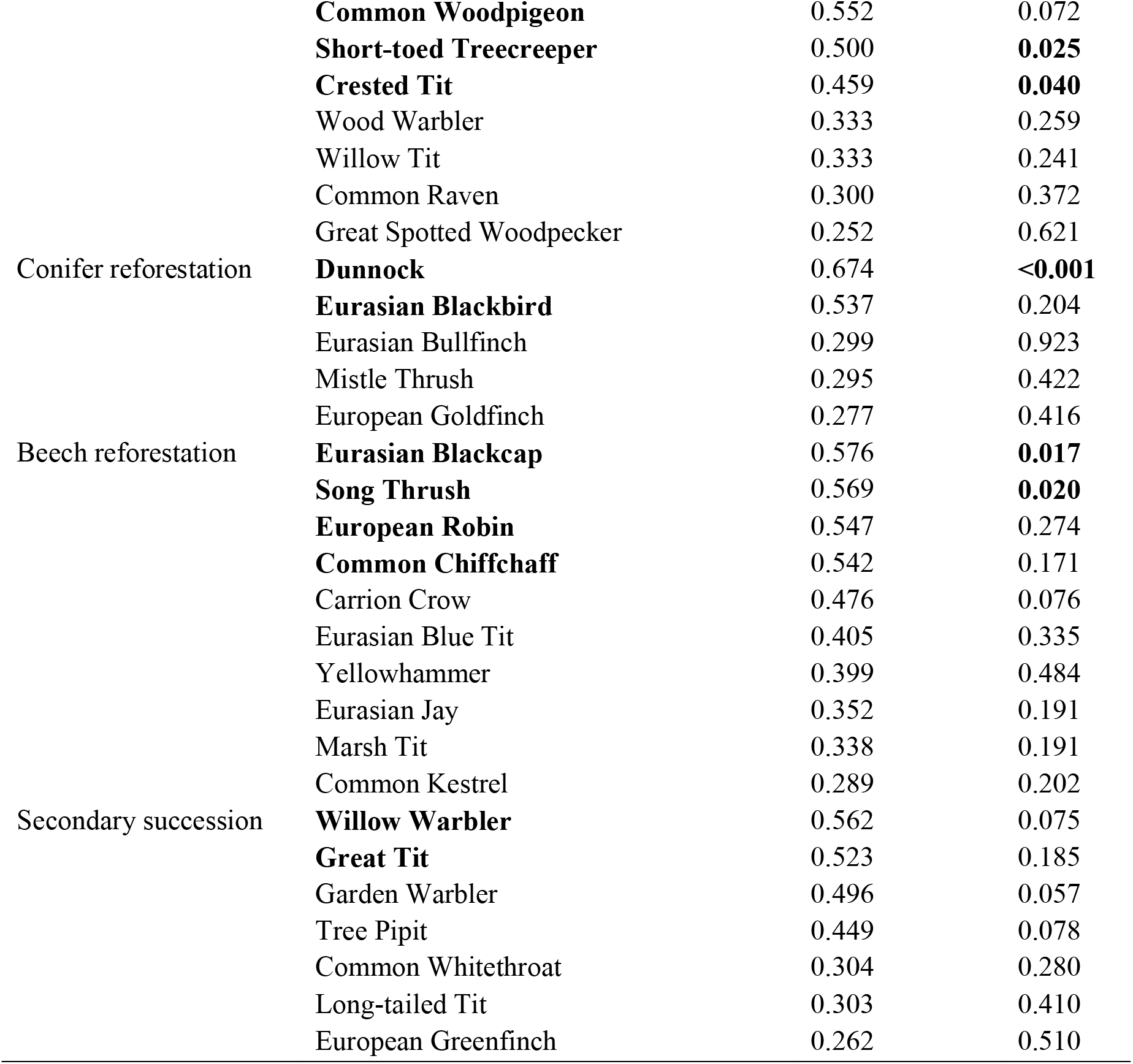
Indicator species with indicator value (IV) ≥ 0.25, separately per habitat and reforestation category. Species with high indicator values (IC > 0.5) and those with significant values (p > 0.05) are typeset in bold. For a full set of recorded species, see the supporting online material, Table S2.

### Community composition

The NMDS results suggested considerable dissimilarities in the community composition of old spruce forest, reforestations and secondary succession. Partitioned along the first ordination axis, stands of old spruce forest supported very different bird communities from those of reforestations or secondary succession, which resembled each other more. Reforestation and secondary succession (arranged on the second NMDS axis) were generally rather similar in community composition, but plots in conifer reforestations were structurally more similar than those in beech reforestations that showed more variation across plots, especially in terms of pioneer tree cover (Fig. 4).

**Figure 4:**
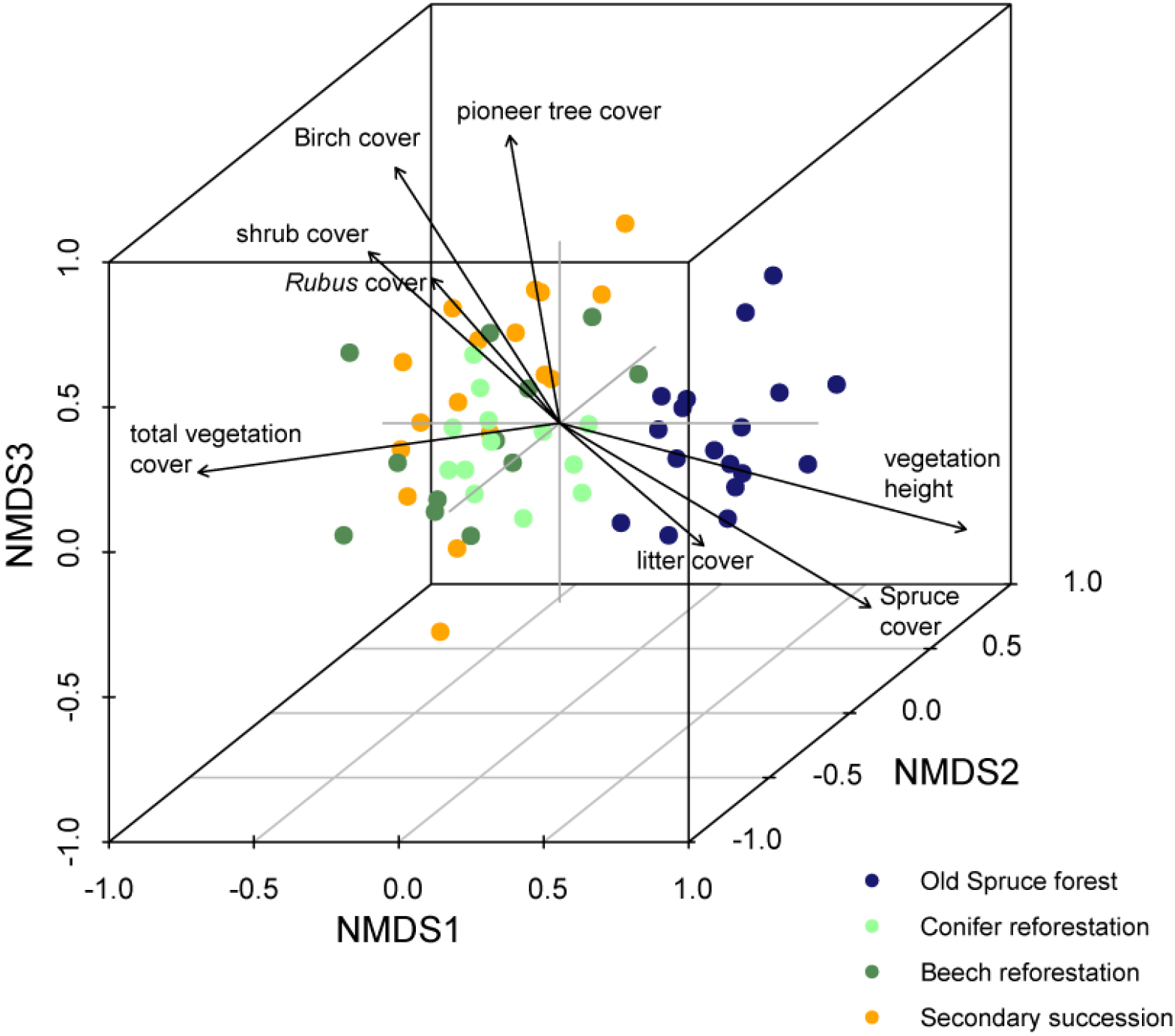
Three-dimensional non-metric multidimensional scaling (NMDS) of bird communities on 60 plots, differentiated by habitat type, with a stress value of 0.168. Plots are displayed as coloured large dots. Black arrows indicate highly significant environmental variables (p<0.001).

### Trait-based analyses

Tendencies for abundances of functional groups to vary between habitat types were visible in nearly all traits examined (Fig. 5), with a number of significant differences (supplementary online material, Table S4). Long-distance migrants were predicted to have significantly higher numbers (p < 0.001) in secondary succession and reforestation compared to the original old spruce forest. Resident species showed an indifferent pattern, whereas short-distance migrants were clearly losers of the disturbance event, and reached comparable abundances to old spruce forest only in conifer reforestations, with significantly lower abundances in all other habitat types. Open-cup nesters were significantly less common in old spruce forest compared to conifer reforestation and secondary succession (p < 0.01). Differences in the abundance of dietary groups were not pronounced, and not significant. The same was true for feeding niche, although there was a tendency for trunk feeders to be more abundant in old spruce forest (p < 0.01, but based on five species only).

**Figure 5:**
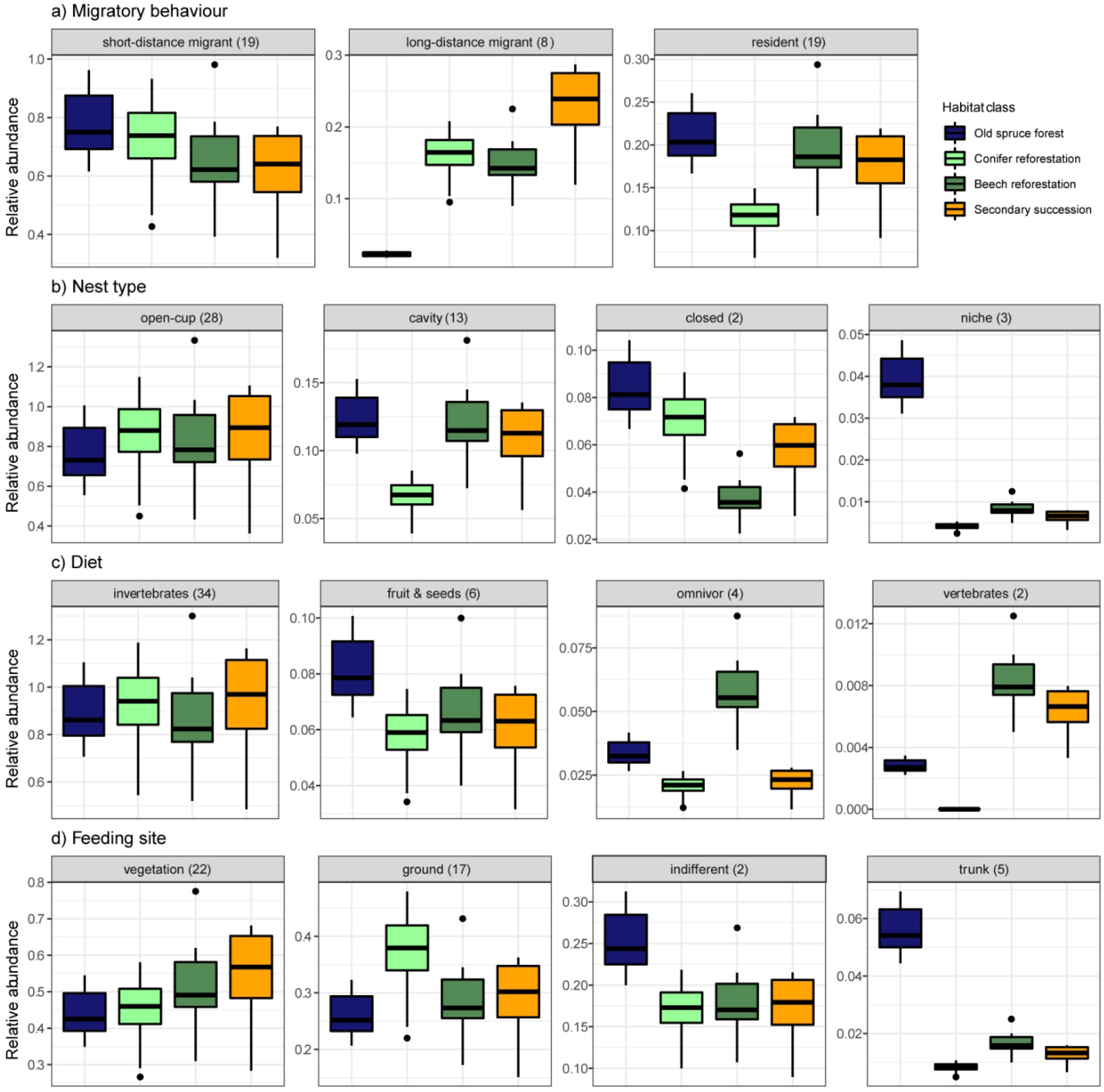
Guild-based plots of predicted relative abundances (from GLMs) for the four different habitat types.

## Discussion

In one of Europe’s largest windthrow landscapes of the past decades, we found that reforestation strategies varied during the first decade after disturbance. In our 1000 ha study area, replanting of non-native conifers predominated with twice the extent of beech reforestation. Large areas were still dominated by successional tree species a decade after the storm, especially birch. Birds responded strongly to windthrow, with a pronounced community turnover. Species associated with high conifer stands reached significantly lower densities at sample plots in disturbed areas. Secondary succession in logged areas and replanted areas was characterized by high densities of long-distance migrants (often species of conservation concern) and shrubland species.

### Reforestation strategies

Across our large study area in the epicentre of windstorm “Kyrill”, all windthrows had been salvage-logged and cleared within two years after the storm, and no areas were left untouched for natural secondary succession as in other large windthrow areas in Europe (Żmihorski & Durska 2011), or in protected areas (Thorn et al. 2016). After clearance, reforestation with conifers (mostly Norway Spruce and Douglas Fir) and beech had happened over large areas, but equally large areas were still dominated by a mix of successional tree species, notably birch. This suggests that local forestry partly considered management recommendations that were developed on a sub-national level after the windstorm (Landesbetrieb Wald & Holz 2007). These recommendations aimed to replace allochthonous fir plantations by mixed stands in the long term, to better adapt forests to future climate-change effects such as drought, windthrow and bark beetle outbreaks. Main recommendations comprised the use of autochthonous tree species (i.e., no conifers) for reforestation, allow natural succession by birch and softwood on cleared areas, and re-plant under the canopy of the emerging pioneer forests. This was assumed to decrease the exposition of “desired”, marketable tree species to wind, sun, radiation and frost, improve the water balance and stop erosion on steep hillslopes (Landesbetrieb Wald & Holz 2007). Economic benefits were suggested to arise from using wood resulting from thinning of these pioneer forests, and by a reduced number of seedlings needed for reforestation (Landesbetrieb Wald & Holz 2007). Not only the windthrow event itself, but also the consideration of the recommended reforestation strategies resulted in a very high diversity of reforestation approaches. Future forests in the area will therefore be more diverse, contain more tree species and be closer to a hypothesized “natural vegetation” than at any point during the past 100 years, when uniform, dark and species-poor spruce plantations dominated vast areas.

### Bird responses to windthrow and reforestation

No differences in species richness and diversity were detected when comparing old spruce forest with reforested areas, and within the reforested areas. This suggests that the loss of old-growth, conifer-attached bird species was offset by the establishment of open-habitat species a decade after the disturbance event, irrespectively of reforestation strategy. This finding is at variance with other studies: in Scots Pine forests in Poland, disturbed sites had a higher species richness and diversity five years after disturbance than undisturbed forest (Żmihorski 2010), whereas in undisturbed Norway Spruce stands in Southern Germany held more species than windthrow area three to seven years after disturbance (Thorn et al. 2016). However, patterns in species richness and diversity are difficult to compare across study sites without prior standardization, partly because of the varying environmental and biogeographical conditions that will results in different baseline species richness. The loss of old spruce could also lead to an increase in the population density at remaining stands as birds from thrown areas resettle there, but we were not able to test this due to the limitations of our space-for-time approach.

Strong differences in abundance and community composition were found when comparing mature, undisturbed forest with disturbance areas. This is unsurprising, as vegetation structure changed from high forest to shrubby vegetation and low, small trees (<10m) due to the disturbance event. Earlier studies on bird responses to windthrow found similar patterns, with an increase of open-habitat and edge-habitat species (Żmihorski 2010, Thorn et al. 2016). Despite different climatic conditions, altitude and tree species composition, the winner and loser species across post-disturbance studies in Europe were surprisingly similar. Losers repeatedly featured Gold- and Firecrest, Coal Tit and Crested Tit, Chaffinch and Treecreepers. Among winners, often Yellowhammer, Tree Pipit, Willow Warbler and Dunnock were named (this study, Thorn et al. 2016, Żmihorski 2010, Żmihorski and Durska 2011).

Our trait-based analysis suggested that open areas hosted many long-distance migrants, whereas undisturbed mature spruce was characterized by significantly more resident species (Thorn et al. 2016). Other traits, such as nest type, diet or feeding site hardly differed between the habitats. The lack of differences might be explained by the rather small habitat patches, and therefore close vicinity of other habitats: Cavity-nesters such as woodpeckers, for example, were observed to use the disturbed and reforested areas for feeding.

Few significant differences in species abundances were found when comparing areas of different reforestation strategy, i.e., areas dominated either by natural succession or by planted conifers or by planted beech. However, the indicator species analysis revealed a number of species (with indicator value >25) for each of the reforested habitat types. Conifer and beech reforestations were characterised by rather common, ubiquitous forest species. Secondary succession was characterized by a number of species of conservation concern that are declining in Germany (Gerlach et al. 2019) and across Europe (Gregory et al. 2019), including the long-distance migrants Willow Warbler, Tree Pipit and Garden Warbler. Our results therefore suggest that the strategy of allowing secondary succession in a considerable part of the study area benefitted depleted species.

### Conclusions

The projected future increase in forest disturbance, including windthrow, might not only cause losses but also be an opportunity for biodiversity in Central Europe. Disturbance disproportionally often affects allochthonous tree species and plantations. We here show, as other authors, that mostly biodiversity associated with these stands, generally unthreatened species will be lost. Furthermore, forests have gradually become darker, cooler and denser during the past 200 years, due to the near-complete abandonment of wood pasture and coppicing (McGrath et al. 2015, Plieninger et al. 2015). More recently, the implementation of policies avoiding clear-cuts and promoting continuous tree cover (Thorn et al. 2019), as well as a decrease in demand for firewood and increased wood volume due to raised atmospheric CO_2_-levels have resulted in a further darkening of the forests. This has led to the decline of light-demanding plant and animal species (incl. long-distance migratory birds) that were adapted to frequent natural and anthropogenic forest disturbance (Thorn et al. 2019). For example, for 16 % of the butterfly species occurring in Germany (n = 187) a loss of traditional woodland management early successional stages is considered a major threat (Reinhardt et al. 2020). Man-made clearcuts can harbour significant populations of threatened farmland birds and butterflies (Ram et al. 2020). Whether new opportunities of large-scale forest disturbance can be harnessed by open-country species will depend on species-specific traits such as dispersal ability, but also on the future management of forest disturbance areas (Müller et al. 2019). For economic reasons, strategies that allow natural succession in disturbed areas, eventually resulting in a near-natural tree composition, are rarely applied. Usually, salvage logging takes place, followed by a complete clearance of the area and fast reforestation with either native or introduced tree species, as in our study. Salvage logging has been shown to significantly change species communities, decrease the richness of saproxylic organisms, but increase the richness of non-saproxylic taxa in comparison with uncleared areas (Thorn et al. 2017). We here show that even in salvage-logged areas, attempts to diversify post-disturbance regrowth, and allowing a certain cover of secondary succession will result in gains in declining species.

## Supporting information

TextS1_mapping_results

TableS1_bird_data

TableS2_trait_data

TableS3_distance_sampling_results

TableS4_GLM_results

## Acknowledgements

We thank Richard Nikodem of Forestry department Balve for an introduction to the study area, indispensable help with forest inventory data and maps, and permits to access the study area. We also thank the hunting associations and private forest owners for allowing access to their land. This research was unfunded.

## CRediT author statement

*Johannes Kamp*: Conceptualization, Methodology, Formal Analysis, Writing – Original Draft, Supervision; *Johanna Trappe*: Methodology, Sofware, Formal Analysis, Writing – Original Draft, *Luca Dübbers*; Investigation, Formal Analysis, Writing – Review and editing, *Stephanie Funke*: Investigation, Formal Analysis, Writing – Review and editing.

## References

Allen, C.D., et al., 2010. A global overview of drought and heat-induced tree mortality reveals emerging climate change risks for forests. For. Ecol. Manage. 259, 660–684.

Bouget, C., Duelli, P., 2004. The effects of windthrow on forest insect communities: a literature review. Biol. Cons. 118, 281–299.

Buckland, S.T., Rexstad, E., Marques, T.A., Oedekove, C.S., 2015. Distance sampling: Methods and applications. Springer, New York.

Burghoff, O., Hauner, O., Kubik, A., 2010. Weather-driven natural hazards - Approaches of risk management in the German insurance industry. Geophysical Research Abstracts 12, EGU2010–15081

De Caceres, M., P. Legendre, 2009. Associations between species and groups of sites: indices and statistical inference. Ecology 90, 3566–3574.

Dobor, L., Hlásny, T., Rammer, W., Zimová, S., Barka, I., Seidl, R., 2020. Spatial configuration matters when removing windfelled trees to manage bark beetle disturbances in Central European forest landscapes. J. Environm. Manage. 254, 109792.

Federal Ministry of Food and Agriculture (2015). The forests in Germany. Selected results of the third national forest inventory. Download: https://www.bmel.de/SharedDocs/Downloads/EN/Publications/ForestsInGermany-BWI.pdf?_blob=publicationFile

Fink, A. H., Brücher, T., Ermert, V., Krüger, A., Pinto, J. G., 2009. The European storm Kyrill in January 2007: synoptic evolution, meteorological impacts and some considerations with respect to climate change. Nat. Haz. Earth Sys. Sci. 9, 405–423.

Fiske, I., Chandler, R., 2011. Unmarked: an R package for fitting hierarchical models of wildlife occurrence and abundance. J. Stat. Soft. 43, 1–23.

Fukui, D., Hirao, T., Murakami, M., Hirakawa, H., 2011. Effects of treefall gaps created by windthrow on bat assemblages in a temperate forest. For. Ecol. Manage. 261, 1546–1552.

Georgiev, K. B., Chao, A., Castro, J., Chen, Y. H., Choi, C. Y., Fontaine, J. B., Hutto, R.L., Lee, E.-J., Müller, J., Żmihorski, M., Thorn, S., 2020 (in press). Salvage logging changes the taxonomic, phylogenetic and functional successional trajectories of forest bird communities. Journal of Applied Ecology, doi: 10.1111/1365-2664.13599

Gerlach, B., Dröschmeister, R., Langgemach, K., Borkenhagen, K., Busch, M., Hauswirth, M., Heinecke, T., Kamp, J., Karthäuser, J., König, C., Markones, N., Prior, N., Trautmann, S., Wahl, J., Sudfeldt, C., 2019. Birds in Germany – an overview of trends and population sizes. DDA, BfN, LAG VSW, Münster. Download: https://www.bfn.de/fileadmin/BfN/monitoring/Dokumente/ViD_Uebersichten_zur_Bestandssituation.pdf [In German.]

Gregory, R. D., Skorpilova, J., Vorisek, P., Butler, S., 2019. An analysis of trends, uncertainty and species selection shows contrasting trends of widespread forest and farmland birds in Europe. Ecol. Ind. 103, 676–687.

Gregow, H., Laaksonen, A., Alper, M. E., 2017. Increasing large scale windstorm damage in Western, Central and Northern European forests, 1951–2010. Sci. Reports 7, 46397.

Handbook of the Birds of the World and BirdLife International, 2019. Digital checklist of the birds of the world. Version 4. Download: http://datazone.birdlife.org/userfiles/file/Species/Taxonomy/HBW-BirdLife_Checklist_v4_Dec19.zip

Hartig, F., 2019. DHARMa: Residual Diagnostics for Hierarchical (Multi-Level / Mixed) Regression Models. R package version 0.2.4. Download : https://CRAN.R-project.org/package=DHARMa

Ishizuka, M., Ochiai, Y., Utsugi, H., 2002. Microenvironments and growth in gaps. Pp. 229–244 in: Nakashizuka, M. (Ed.), Diversity and Interaction in a Temperate Forest Community: Ogawa Forest Reserve of Japan. Springer, Tokyo,.

Jonášová, M., Vávrová, E., Cudlín, P., 2010. Western Carpathian mountain spruce forest after a windthrow: natural regeneration in cleared and uncleared areas. For. Ecol. Manage 259, 1127–1134.

Kéry, M., Royle, A., 2016. Applied Hierarchical Modeling in Ecology: Analysis of distribution, abundance and species richness in R and BUGS: Volume 1: Prelude and Static Models. Academic Press, London.

Kirby, K., Watkins, C., 2015. Europe’s changing woods and forests: from wildwood to managed landscapes. CABI.

Landesbetrieb Wald & Holz Nordrhein-Westfalen, 2007: Recommendations for the reforestation of windstorm-affected areas in Northrhine-Westphalia. Münster, Germany. Download : https://www.wald-und-holz.nrw.de/fileadmin/Publikationen/Broschueren/Broschuere_Empfehlungen_Wiederbewaldung_Orkanflaechen.pdf [In German.]

Lenth, R., 2019. emmeans: Estimated Marginal Means, aka Least-Squares Means. R package version 1.4.1. Download : https://CRAN.R-project.org/package=emmeans

Leverkus, A. B., Lindenmayer, D. B., Thorn, S., Gustafsson, L., 2018. Salvage logging in the world’s forests: Interactions between natural disturbance and logging need recognition. Global Ecol. Biogeo. 27, 1140–1154.

Lindner, M., Maroschek, M., Netherer, S., Kremer, A., Barbati, A., Garcia-Gonzalo, J., Seidl, R., Delzon, S., Corona, P., Kolström, M., Lexer, M., Marchetti, M., 2010. Climate change impacts, adaptive capacity, and vulnerability of European forest ecosystems. For. Ecol. Manage. 259, 698–709.

McDowell, N. G., Allen, C., Anderson-Teixeira, K., Aukema, B.H., Bond-Lamerty, B., Chini, L., Clark, J.S., Dietze, M., Grossiord, C., Hanbury-Brown, A., Hurtt, G.C., Jackson, R.B., Johnson, D.J., Kueppers, L., Lichstein, J.W., Ogle, K., Poulter, B., Pugh, T.A.M., Seidl, R., Turner, M., Uriarte, M., Walker, A.P., Xu, C., 2020. Pervasive shifts in forest dynamics in a changing world. Science 368, 6494.

McGrath, M. J., Luyssaert, S., Meyfroidt, P., Kaplan, J. O., Burgi, M., Chen, Y., Erb, K., Gimmi, U., McInerney, D., Naudts, K., Otto, J., Pasztor, F., Ryder, J., Schelhaas, M.-J., Valade, A., 2015. Reconstructing European forest management from 1600 to 2010. Biogeosciences 12, 4291–4316.

Mehr, M., Brandl, R., Kneib, T., Müller, J., 2012. The effect of bark beetle infestation and salvage logging on bat activity in a national park. Biodiv. Cons. 21, 2775–2786.

Müller, J., Noss, R. F., Thorn, S., Bässler, C., Leverkus, A. B. Lindenmayer, D., 2019. Increasing disturbance demands new policies to conserve intact forest. Cons. Lett. 12, e12449.

Murakami, M., Hirao, T., Iwamoto, J., Oguma, H., 2008. Effects of windthrow disturbance on a forest bird community depend on spatial scale. Basic Appl. Ecol. 9, 762–770.

Nachtergale, L., Ghekiere, K., De Schrijver, A., Muys, B., Luyssaert, S., Lust, N., 2002. Earthworm biomass and species diversity in windthrow sites of a temperate lowland forest. Pedobiologia 46, 440–451.

Oksanen, J., Blanchet, F. G., Friendly, M., Kindt, R., Legendre, P., McGlinn, D., Minchin,P. R., O’Hara, R. B., Simpson, G. L., Solymos, P., Stevens, M. H. H, Szoecs, E. & H. Wagner, 2018. vegan: Community Ecology Package. R package version 2.5-2. Download : https://CRAN.R-project.org/package=vegan

Opengeodata NRW (2020). Kyrill windthrow areas in Northrhine-Westfalia, shapefile. Download: https://www.opengeodata.nrw.de/produkte/umwelt_klima/wald_forst/wald/windwurfschadflaechen-kyrill_EPSG25832_Shape.zip

Panferov, O., Doering, C., Rauch, E., Sogachev, A., Ahrends, B., 2009. Feedbacks of windthrow for Norway spruce and Scots pine stands under changing climate. Envir. Res. Lett. 4, 045019.

Plieninger, T., Hartel, T., Martín-López, B., Beaufoy, G., Bergmeier, E., Kirby, K., Montero, M.J., Moreno, G., Oteros-Rozas, E., Van Uytvanck, J. 2015. Wood-pastures of Europe: Geographic coverage, social– ecological values, conservation management, and policy implications. Biol. Cons. 190, 70–79.

R Core Team (2020). R: A language and environment for statistical computing. R Foundation for Statistical Computing, Vienna, Austria. Download : https://www.R-project.org/.

Rahmstorf, S., Coumou, D., 2011. Increase of extreme events in a warming world. Proc. Nat. Acad. Sci. Am. 108, 17905–17909.

Ram, D., Lindström, Å., Petterson, L.A., Caplat, P., 2020. Forest clearcuts as habitat for farmland birds and butterflies. For. Ecol. Manage. 473: 118239.

Reinhardt, R., Harpke, A., Caspari, S., Dolek, M., Kühn, E., Musche, M., Trusch, R., Wiemers, M., Settele, J. (2020). Distribution atlas of the butterflies and burnet moths of Germany. Ulmer, Stuttgart. [In German.]

Ruel, J. C., 1995. Understanding windthrow: silvicultural implications. For. Chronicle 71, 434–445.

Schelhaas, M.-J., Nabuurs, G.-J., Schuck, A., 2003. Natural disturbances in the European forests in the 19th and 20th centuries. Glob. Change Biol. 9:1620–163.

Schuldt, B., Buras, A., Arend, M., Vitasse, Y., Beierkuhnlein, C., Damm, A., Gharun, M., Grams, T.E.E., Hauck, M., Hajek, P., Hartmann, H., Hiltbrunner, E., Hoch, G., Holloway-Phillips, M., Körner, C., Larysch, E., Lübbe, R., Nelson, D.E., Rammig, A., Rigling, A., Rose, L., Ruehr, N.K., Schumann, K., Weiser, F., Werner, C., Wohlgemuth, T., Zang, C.S., Kahmen, A., 2020 (in press). A first assessment of the impact of the extreme 2018 summer drought on Central European forests. Basic Appl. Ecol., doi : 10.1016/j.baae.2020.04.003

Seidl, R., Schelhaas, M.-J., Lexer, M.J., 2011. Unraveling the drivers of intensifying forest disturbance regimes in Europe. Glob. Change Biol. 17, 2842–2852.

Seidl, R., Thom, D., Kautz, M., Martin-Benito, D., Peltoniemi, M., Vacchiano, G., Wild, J., Ascoli, D., Petr, M., Honkaniemi, J., Lexer, M.J., Trotsiuk, V., Mairota, P., Svoboda, M., Fabrika, M., Nagel, T.A :, Reyer, C.P.O., 2017. Forest disturbances under climate change. Nat. Clim. Change 7, 395–402.

Senf, C., Pflugmacher, D., Zhiqiang, Y., Sebald, J., Knorn, J., Neumann, M., Hostert, P., Seidl, R., 2018. Canopy mortality has doubled in Europe’s temperate forests over the last three decades. Nat. Comm. 9, 4978.

Senf, C., Sebald, J., Seidl, R., 2020. Increasing canopy mortality challenges the future of Europe’s forests. bioRxiv preprint, doi:10.1101/2020.03.30.015818

Senf, C., Seidl, R., 2020. Mapping the coupled human and natural disturbance regimes of Europe’s forests. bioRxiv preprint, doi: 10.1101/2020.03.30.015875

Steinkamp, J.,, Hickler, T., 2015. Is drought-induced forest dieback globally increasing? J. Ecol. 103, 31–43.

Thorn, S., Werner, S. A., Wohlfahrt, J., Bässler, C., Seibold, S., Quillfeldt, P., Müller, J., 2016. Response of bird assemblages to windstorm and salvage logging – Insights from analyses of functional guild and indicator species. Ecol. Ind. 65, 142–148.

Thorn, S., Bässler, C., Brandl, R., Burton, P. J., Cahall, R., Campbell, J. L., Castro, J., Choi, C.-Y., Cobb, T., Donato, D.C., Durska, E., Fontaine, J.B., Gauthier, S., Hebert, C., Hothorn, T., Hutto, R.L., Lee, E.-J., Leverkus, A.B., Lindenmayer, D.B., Obrist, M.K., Rost, J., Seibold, S., Seidl, R., Thom, D., Waldron, K., Wermelinger, B., Winter, M.-B., Żmihorski, M., Müller, J., 2018. Impacts of salvage logging on biodiversity: A meta-analysis. J. Appl. Ecol. 55, 279–289.

Thorn, S., Müller, J., Leverkus, A. B., 2019. Preventing European forest diebacks. Science 365, 1388.

Trenberth, K. E., Fasullo, J. T., Shepherd, T. G., 2015. Attribution of climate extreme events. Nat. Clim. Change 5, 725–730.

von Oheimb, G., Friedel, A., Bertsch, A., Härdtle, W., 2007. The effects of windthrow on plant species richness in a Central European beech forest. Plant Ecology 191, 47–65.

Wald & Holz NRW, 2019. Windstorm Kyrill and its consequences. Download: https://www.wald-und-holz.nrw.de/wald-in-nrw/wald-und-klima/kyrill-und-seine-folgen-in-nrw [In German.]

Ulanova, N. G., 2000. The effects of windthrow on forests at different spatial scales: a review. For. Ecol. Manage. 135, 155–167.

Żmihorski, M., 2010. The effect of windthrow and its management on breeding bird communities in a managed forest. Biodiv. Cons. 19, 1871–1882.

Żmihorski, M., Durska, E., 2011. The effect of contrasting management types on two distinct taxonomic groups in a large-scaled windthrow. Europ. J. Forest Res. 130, 589–600.

