## Supplementary material for "Impacts of landscape-scale windthrow and subsequent, variable reforestation on bird communities in Central Europe": TextS1_mapping_results

Fig. S1: Screenshot (700 x 400 m) of an aerial image (ortho-rectified, 0.4 m resolution) from google earth used to map forest disturbance, taken on 31^st^ March 2009. Numbers mark example areas to illustrate the mapping approach (cf. main text): 1 –spruce, not thrown (partly salvage-logged in lower right corner), 2 – beech, not thrown, 3, 4, 6, 7 and 8 – spruce windthrow, 5 – beech windthrow (note irregularly
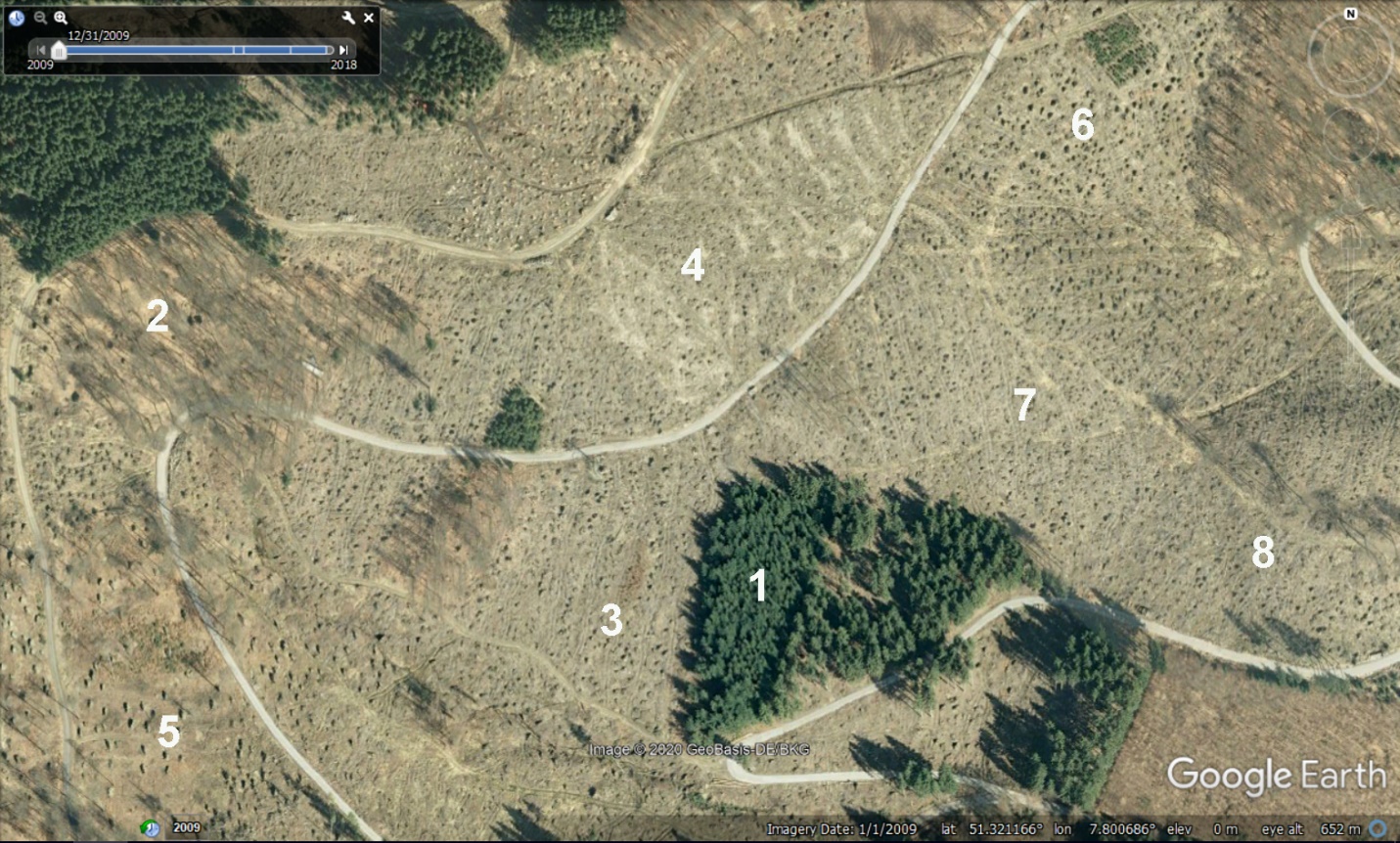
distributed, large uprooted trunk bases and browner soil surface colour compared to spruce windthrow).

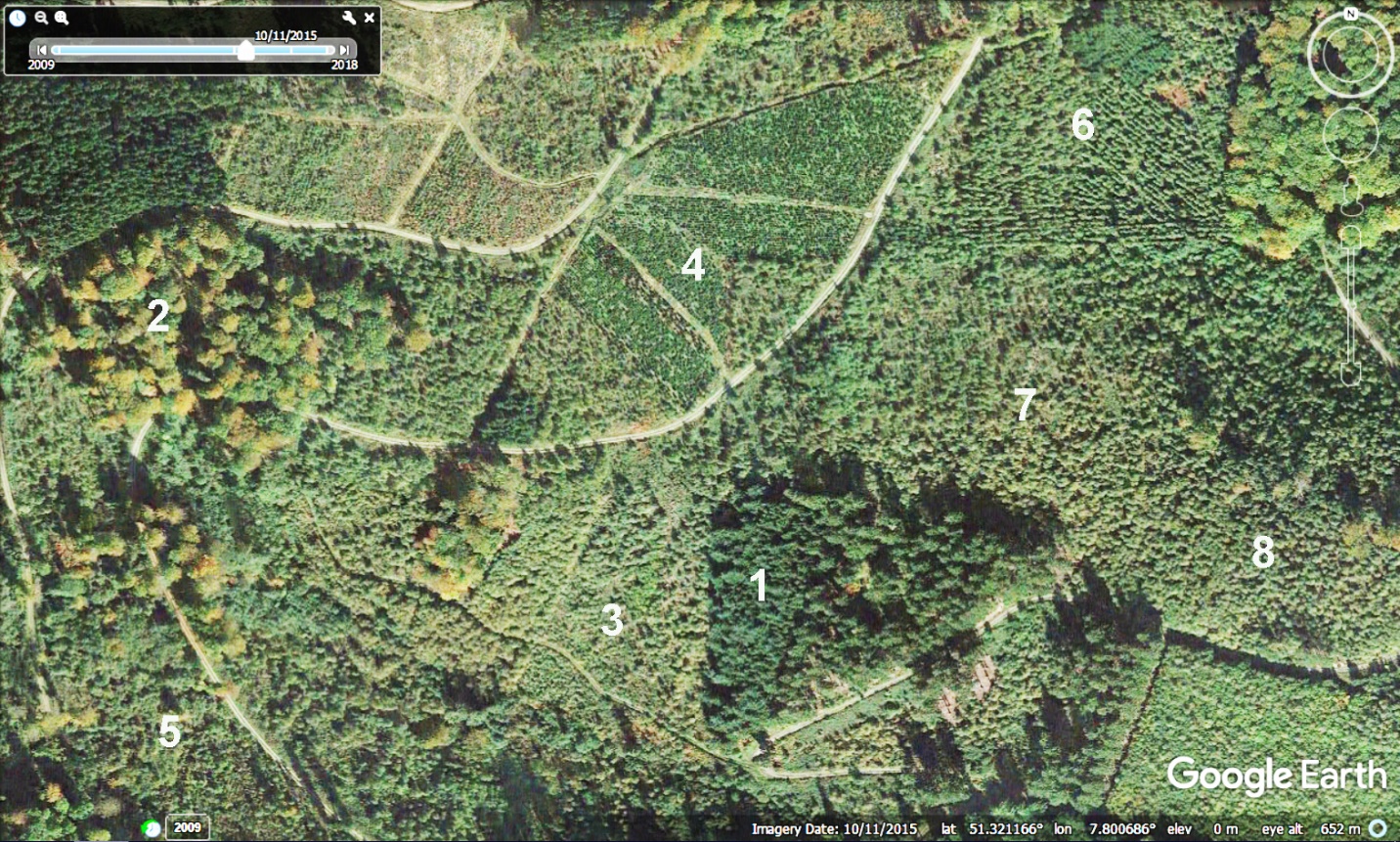
Fig. S2: Screenshot (700 x 400 m) of the same area as in Fig. S1, aerial image (ortho-rectified, 0.4 m resolution) from google earth taken on 11th October 2015. Numbers mark example areas to illustrate the mapping approach (cf. main text): 1 –spruce, not thrown (partly salvage-logged in lower right corner), 2 – beech, not thrown (rusty leaf colouration in autumn), 3, 5, 7 and 8 – not yet reforested windthrows, 4 –conifer reforestation (darker green colour, regularly spaced), 6 – larch reforestation (pale green, regularly spaced).

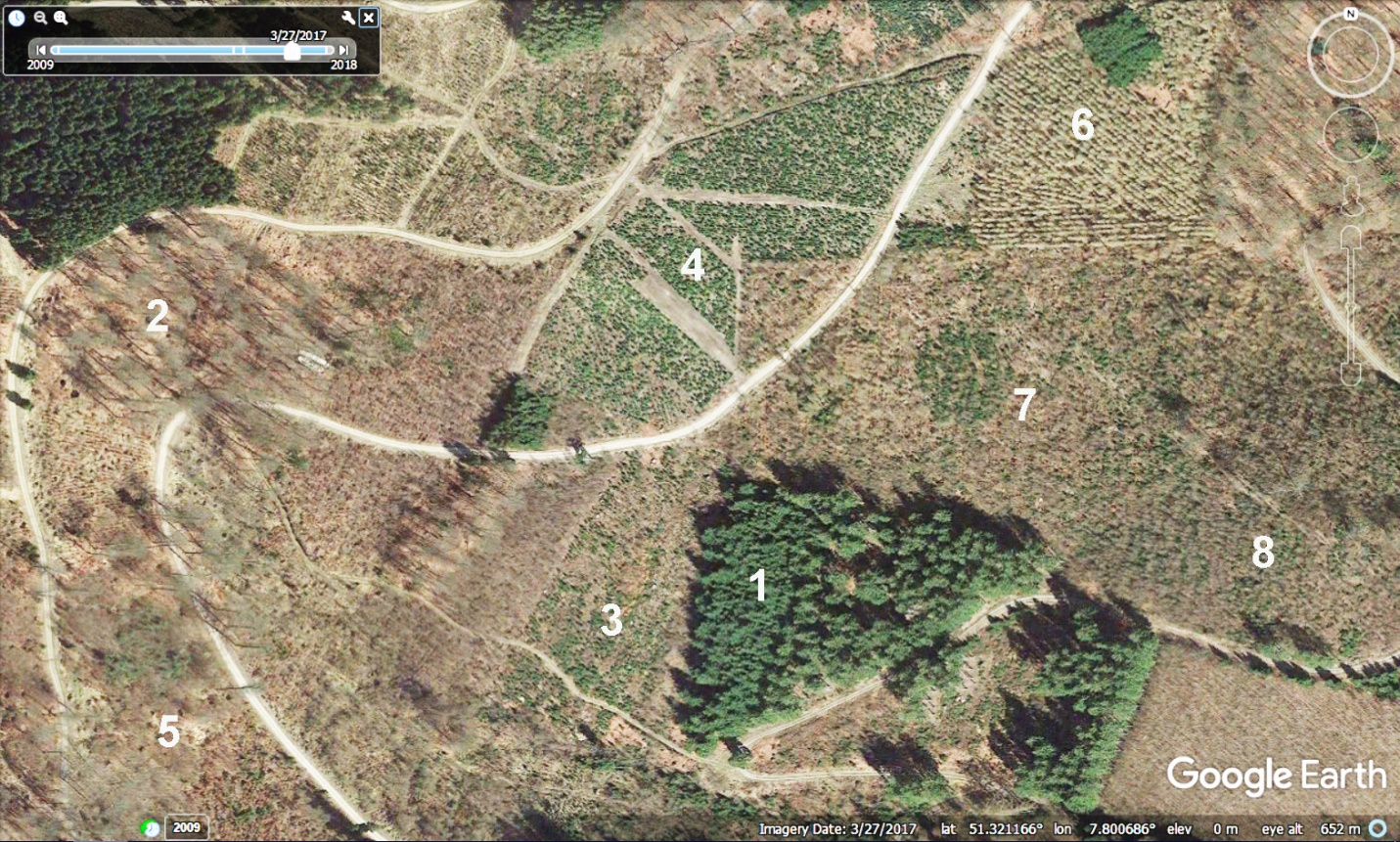
Fig. S3: Screenshot (700 x 400 m) of the same area as in Fig. S1, aerial image (ortho-rectified, 0.4 m resolution) from google earth taken on 27th March 2017. Numbers mark example areas to illustrate the mapping approach (cf. main text): 1 –spruce, not thrown (partly salvage-logged in lower right corner), 2 – beech, not thrown, 3 – spruce plantation, 4 – Douglas fir plantation (for Christmas trees, with tracks for cutting access), 5 – not reforested beech regrowth, 6 – larch reforestation (pale brown, regularly spaced), 7 – natural regrowth (deciduous trees dominating, “combed” pattern in upper part suggests replanting of beech in the understory without prior surface treatment), 8 – natural regrowth, not replanted.

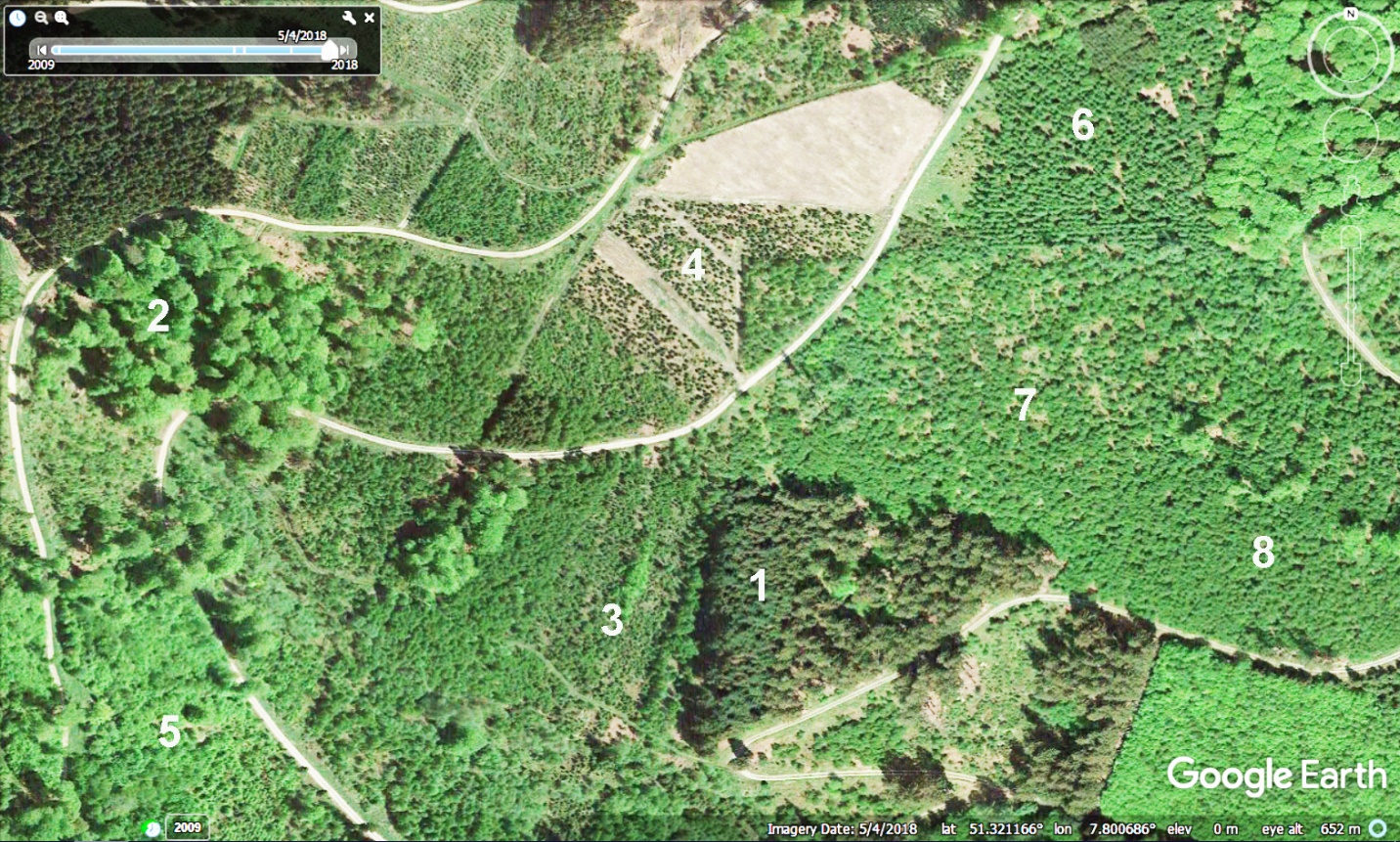
Fig. S4: Screenshot (700 x 400 m) of the same area as in Fig. S1, aerial image (ortho-rectified, 0.4 m resolution) from google earth taken on 4th May 2018. Numbers mark example areas to illustrate the mapping approach (cf. main text) as in Fig. S3, but part of 4 (conifer plantation) now harvested for sale as Christmas trees, deciduous trees in 7 and 8 (dark green) now distinguishable as birch from pale green beech regrowth (5).
